# EasyFigAssembler: Enhance Omics Data Storytelling through Effective Figure Assembly

**DOI:** 10.1101/2025.10.01.679913

**Authors:** Nguyen Phuoc Long, Nguyen Quang Thu

## Abstract

**Motivation:** An effective figure helps communicate the data and findings clearly. The assembly of multi-panel omics-based figures for scientific publications is a time-consuming and error-prone process that often requires specialized software and skills. However, there is a lack of a quick and effective tool for assembling complex omics-based figures.

**Results:** We present EasyFigAssembler, a user-friendly web application that significantly simplifies and accelerates the making of multi-panel omics-based figures. The app supports non-destructive panel editing, multi-figure project management, and high-resolution export to TIFF, PDF, PNG, and JPEG formats, providing a complete solution for figure assembly without requiring programming expertise. Its Smart Layout feature automates the initial arrangement of panels into optimal, publication-ready grids. The advanced controls for column and row spanning enable the rapid assembly of complex, non-uniform layouts that are typically time-consuming to create. The workflow is ensured by an integrated Quality Advisor module that provides real-time feedback on effective resolution and color-blind-friendly palettes to ensure journal compliance. Finally, users can manage multiple figures and save their projects effortlessly.

**Availability and implementation:** The source code of EasyFigAssembler is available at https://github.com/Pharmaco-OmicsLab/EasyFigAssembler. The web interface for users can be accessed at https://easyverse.app/easyfig/.

## 1. Introduction

Manuscript preparation and formatting prior to journal submission is a tedious and time-consuming step in the biomedical sciences (1). There have been advocates to simplify the requirements of manuscript preparation and formatting to reduce time wasted on this step and speed up the scientific communication (2, 3). Regardless of the requirements for submission, assembling individual image panels into a final complete figure is a key, recurring step in manuscript preparation that is often time-consuming and prone to errors. Researchers have to position panels manually, add labels, and ensure adherence to various and stringent journal-specific guidelines for dimensions, resolution, and font sizes. However, general graphic editors often lack integrated scientific context and quality control. This requires researchers to manually manage complex constraints, such as Dots Per Inch (DPI) and panel alignment, a process that is prone to inconsistency and sub-optimal results.

One of the most commonly used tools for making scientific figures, ggplot2 (4), provides an environment to create and customize multi-panel figures via ggpubr’s facet (5). Besides, patchwork is helpful for combining multiple plots into a single layout (6). However, these tools primarily support figures created within the R environment and cannot easily incorporate external image files, e.g., microscopy TIFFs, pathway diagrams, and network graphs. This hampers the researchers from meeting the standard requirements for publishing high-quality multi-omics research. Programming-based tools also lack intuitive and straightforward customizable functions, which limits their usefulness for a larger user base. Commercial tools offer robust solutions but come with high costs and steep learning curves. Finally, manual assembly in both free and commercial graphic editors suffers from a lack of reproducibility, as the figure cannot be precisely recreated without repeating the entire manual process.

To address these challenges, we developed EasyFigAssembler, a user-friendly, open-source web app designed for assembling omics-based complex figures. EasyFigAssembler significantly simplifies and accelerates the entire figure creation process, from uploading a panel to downloading a publication-ready figure. It provides a “Smart Layout” engine for automated arrangement and an integrated “Quality Advisor” to examine the compliance with journal standards, ensuring a rapid and error-free workflow. The app was created specifically to meet the need for complex figure assembly of typical plots of general omics research. However, it can also serve as a universal platform for figure assembly, benefiting the broad scientific community.

## 2. Materials and methods

### 2.1. System Architecture and Technology

EasyFigAssembler is a web application designed with a client-server architecture to provide an interactive user experience.

#### 2.1.1. Frontend

The user interface was built with standard HTML, CSS, and dynamic JavaScript. All core figure assembly tasks, including layout calculations, panel reordering, and non-destructive editing, are available for the user in the browser. This approach ensures a responsive user experience because adjustments to layout, spacing, and panel properties are rendered immediately on HTML5 without server latency. Non-destructive edits, such as cropping, rotation, brightness adjustments, and annotations, are managed as a set of parameters applied to a source image. The final, edited panel is generated on the fly in the browser for preview and export.

#### 2.1.2. Backend

The backend is a lightweight Python server using the Flask framework (7). High-Resolution Export Processing is an important component of the backend that provides API endpoints for exporting figures into publication-ready formats, including TIFF and PDF. It receives the final canvas rendered as high-resolution PNG data from the user. It uses the Python Pillow library to convert it, correctly embedding the user-specified DPI metadata required by journals. The function panel (on the right side) provides essential journal-specific guidelines (e.g., column widths, minimum font sizes) to the frontend via a JSON API.

### 2.2. User Workflow and Core Features

The user-friendly and intuitive interface of EasyFigAssembler guides users through a logical workflow, from panel import to final export. Users begin by uploading all individual panels for a figure using the Panel Import feature. The interface supports a drag-and-drop area and a standard file selector, with native support for common web formats as well as scientific formats. Upon upload, the “Smart Layout” feature is automatically activated via Layout Arrangement. The module was designed to analyze the number of panels and journal guidelines, arranging them into an optimized grid. This often creates complex layouts with column and row spanning to maximize clarity and resolution. Users can then override this by selecting from predefined grid layouts, such as 2xN, 3xN, or by selecting the “Custom Layout” mode for complete manual control over panel size and position. Panels can be reordered via drag-and-drop on the canvas or in the panel list.

Panel Editing and Annotation is a dedicated editing module for any figure panels that require additional adjustments. This interface provides non-destructive tools for cropping, rotating, and adjusting brightness/contrast. Users can also add annotations such as text, arrows, and rectangles directly onto the panel. Importantly, a live preview can be called to accurately show how the edits will appear in the final figure layout. The “Quality Advisor” module provides real-time guidance to ensure the figure meets publication standards. It is a practical and straightforward quality control feature that ensures proper figure assembly without overwhelming users with unnecessary information. It checks each panel’s effective DPI, warns against color combinations problematic for colorblind individuals, and provides a visual text-size guide for the selected journal. Users can make global adjustments to panel spacing, label styles, and font properties using interactive sidebar controls.

Once the figure is assembled satisfactorily, it can be exported at a user-specified DPI to PDF, TIFF, PNG, or JPEG format. The entire multi-figure project can be saved to a single .json file and reloaded into the application at a later time, preserving all edits, annotations, and settings to ensure reproducibility.

### 2.3. Application Examples

We used EasyFigAssembler for three popular figure assembling scenarios, showcasing its utility for both simple and complex omics-based figure assembly tasks. The examples highlight core attributes of EasyFigAssembler for creating publication-ready omics figures (**Figure 1**). In all use cases, the application’s default settings for panel spacing, label style, and font size served as an excellent baseline suitable for a wide range of journals. Preloaded examples are available and can help users familiarize themselves with the web app in a short period of time.

**Figure 1.**
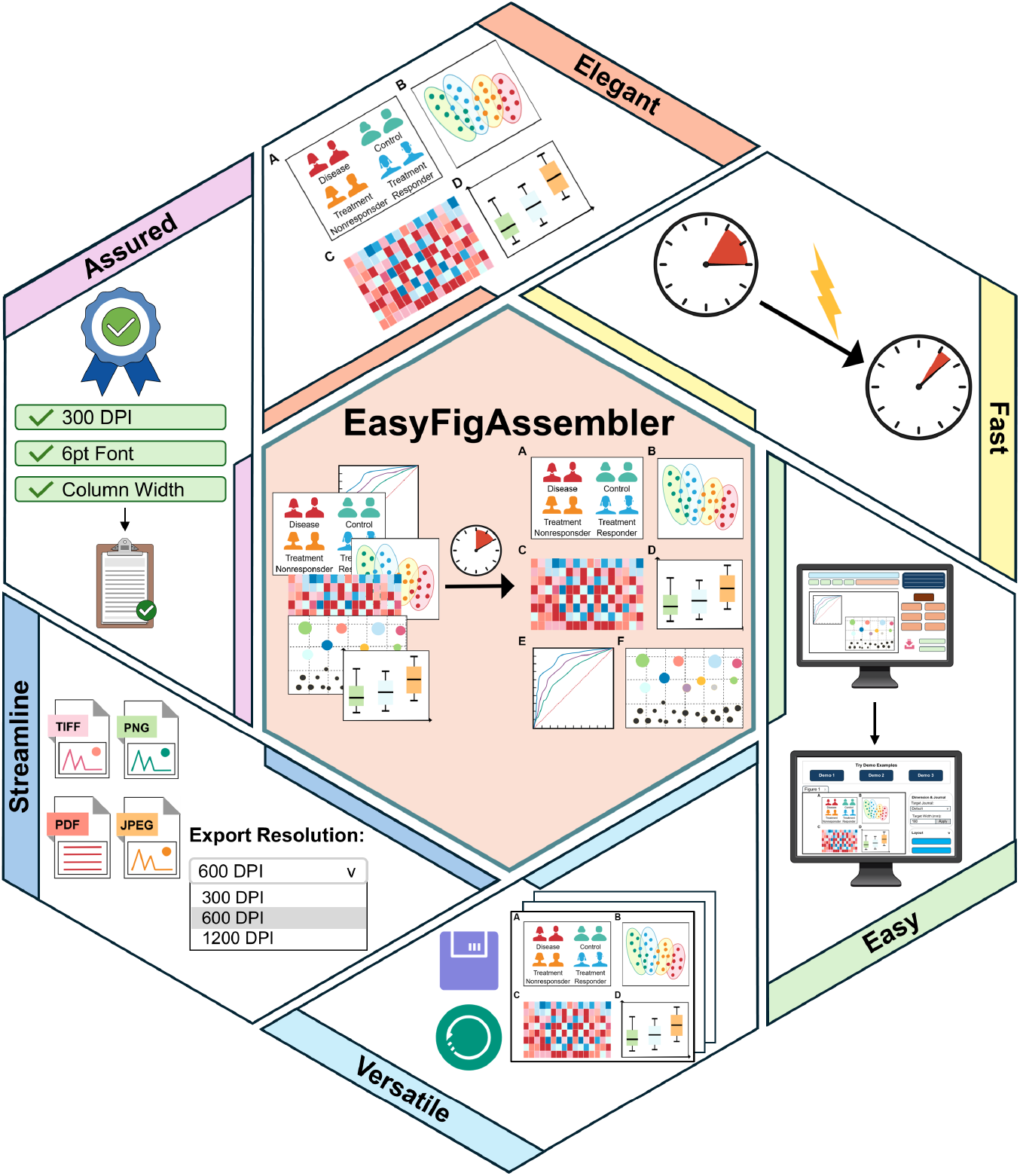
The core attributes of EasyFigAssembler for creating publication-ready scientific figures. This diagram illustrates the six primary attributes of EasyFigAssembler that address the time-consuming and error-prone challenge of manual figure assembly. The application streamlines this process by integrating a Smart Layout engine for automatic arrangement, a Quality Advisor module to ensure journal compliance, and High-Resolution Export options for publication-quality outputs.

#### 2.3.1. Basic Layout with Smart Automation

A standard four-panel figure was assembled using targeted metabolomics data from Feng and colleagues (8), where figure panels were generated and downloaded using MetaboAnalyst 6.0 (9). Upon importing the four panels, the Smart Layout feature automatically generates a publication-ready 2×2 grid. No further manual adjustment is needed in this straightforward use case (**Figure 2**).

**Figure 2.**
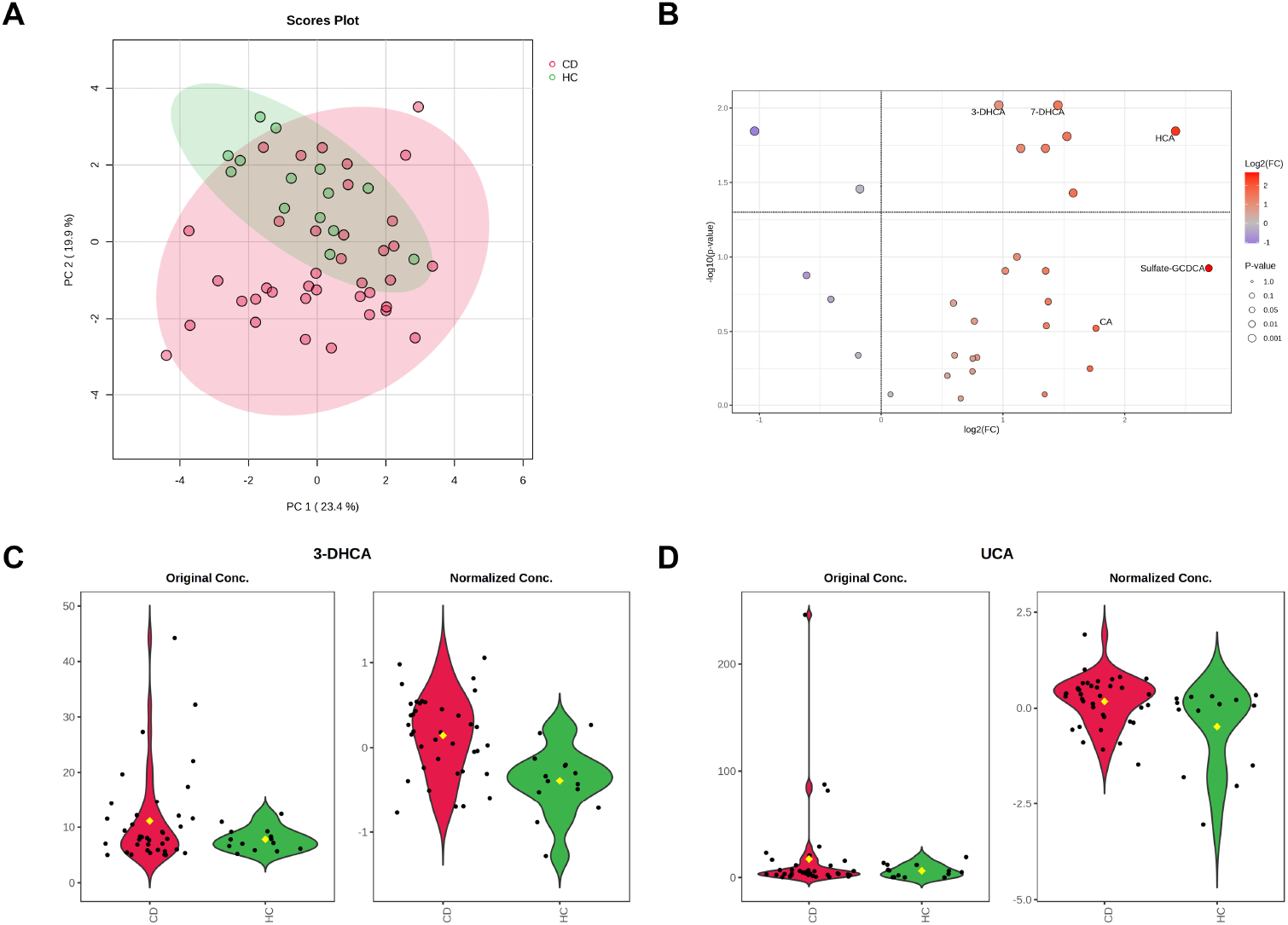
Targeted metabolomics data showcase using the basic layout with automatic arrangement. (A) PCA scores plot. (B) Volcano plot. (C-D) Violin plots. Abbreviations: 3-DHCA: 3-dehydrocholic acid; 7-DHCA: 7-dehydrocholic acid; CA: Cholic acid; CD: Crohn’s Disease; FC: Fold change; HC: Healthy controls; HCA: Hyocholic acid; PCA: Principal component analysis; Sulfate-GCDCA: Glycochenodeoxycholate-3-sulfate; UCA: Ursocholic acid.

#### 2.3.2. Complex Grid with Column Spanning

The fecal microbiome data from the Integrative Human Microbiome Project Consortium, available from MicrobiomeAnalyst 2.0 (10), was used for this example. The generation of a figure from a microbiome dataset required a non-uniform grid to emphasize specific plots. After setting the overall layout to a 3xN grid, we opened the panel edit modal for two of the panels. Their column span was set to “2”, which instantly re-rendered the figure to the desired arrangement (**Figure 3**).

**Figure 3.**
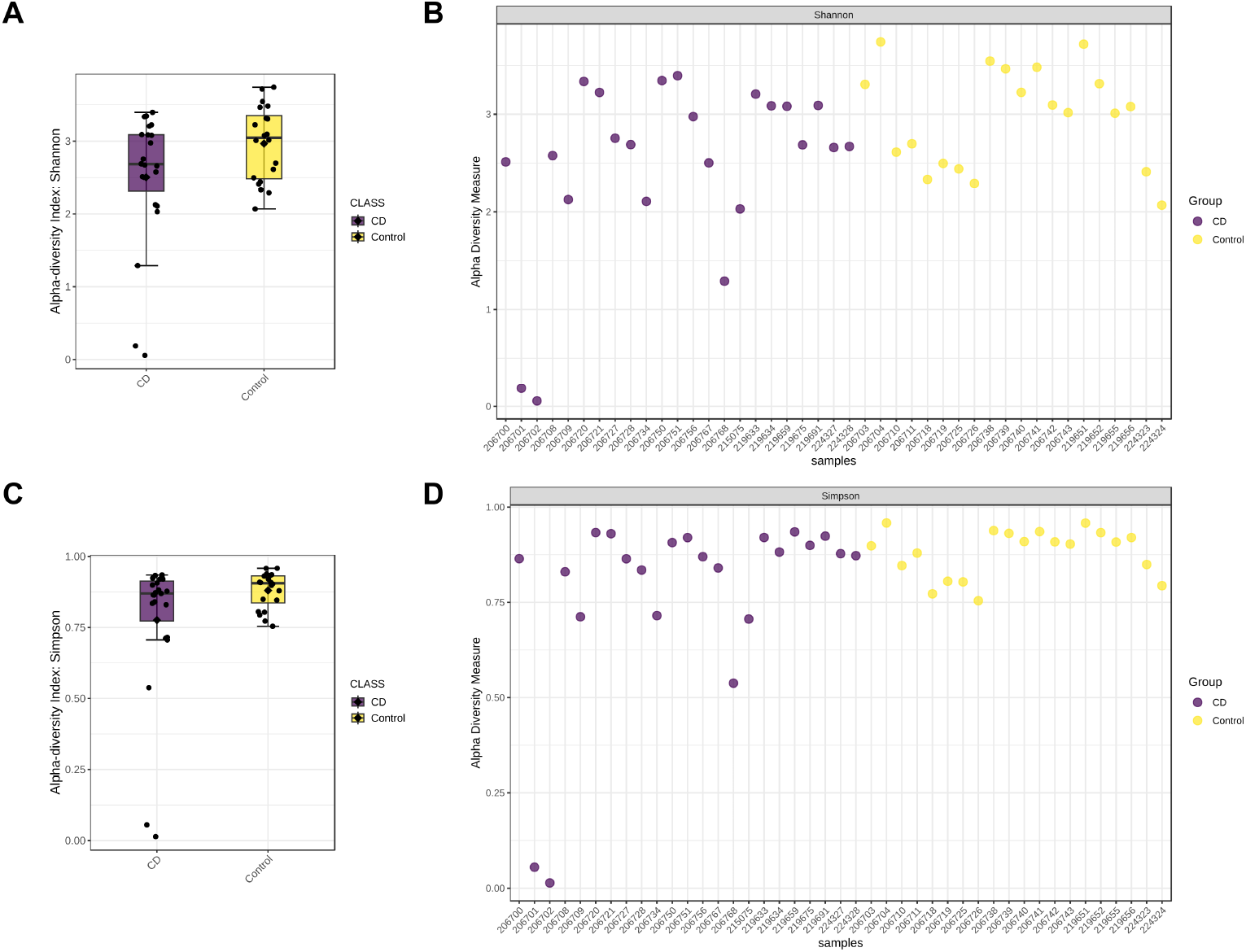
16S microbiome data visualization of alpha-diversity indices using a complex grid with column spanning. (A) and (C). Box plots. (B) and (D). Jitter plots. Abbreviation: CD: Crohn’s disease.

#### 2.3.3. Advanced Multi-Panel Layout

To demonstrate advanced control, a complex seven-panel figure from an untargeted lipidomics study of our group was recreated (11). This required mixed spanning capabilities. The main 2D scores plot was configured to span two columns and two rows (colspan = 2, rowspan = 2). The two heatmap panels were each set to span two columns (colspan = 2). The four box plots remained as 1×1 panels. By applying these settings within each panel’s edit modal and selecting a 4×3 grid, the final, intricate layout was achieved in just a few clicks (**Figure 4**).

**Figure 4.**
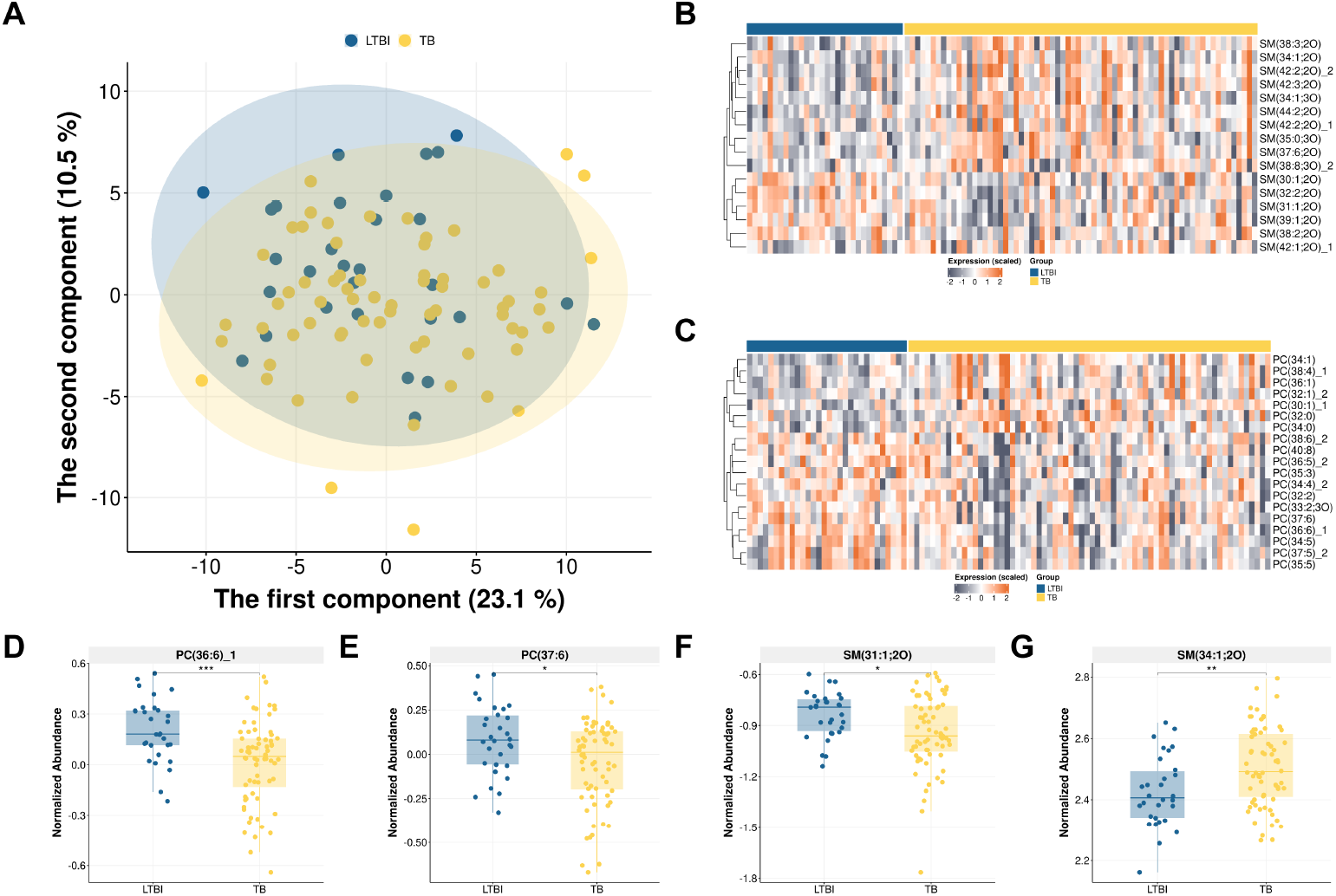
Untargeted lipidomics data visualization with different plot types using a complex grid with various column spanning. (A) PCA scores plot. (B-C) Heatmaps. (D-G). Box plots. Abbreviations: LTBI: Latent Tuberculosis Infection; PC: Phosphatidylcholines; PCA: Principal component analysis; SM: Shingomyelins; TB: Tuberculosis.

## 3. Discussion

Preparing a research proposal or publishing research findings in a peer-reviewed journal can be a time-consuming process (12, 13). In addition to the prolonged period required to properly evaluate a manuscript, the time spent preparing a submission package that adheres to strict journal criteria is also a significant issue (1, 2). In manuscript preparation, creating and assembling figures that accurately represent the narrative of the findings is a tedious and error-prone process. This highlights an unmet need for an app that streamlines the process and reduces the time and effort required by researchers. Here, we developed EasyFigAssembler, an app that provides a straightforward and reproducible workflow for complex figure assembly. An intuitive web-based interface lowers the steep learning curve associated with command-line tools or general-purpose graphics software. Importantly, its integrated “Quality Advisor” provides a layer of insight that traditional tools lack, proactively examining for resolution and color issues to minimize the risk of figures being rejected for technical non-compliance. Furthermore, the non-destructive editing approach and the ability to save and load entire multi-figure projects in a single file are essential for ensuring reproducibility, a fundamental standard in modern science that is challenging to achieve with manual methods.

The application is particularly well-suited for the demands of omics-related research, where publications frequently rely on complex, multi-panel figures that integrate diverse data types, including multiple scores plots of multivariate analysis, heatmaps, volcano plots, pathway diagrams, and network graphs. EasyFigAssembler particularly excels at this by allowing researchers to import various file formats and arrange them using a flexible grid engine that supports mixed column and row spanning. This enables the creation of narrative-driven layouts, allowing scientists to focus on presenting their findings rather than on the technical hurdles of formatting.

The main limitation of this version is that it does not support creating individual figure panels. To complement this, we have developed EasyPubPlot, a tool for generating common plot types used in omics research (14). Using these tools in tandem enables users to create engaging, publication-ready figures while saving valuable research time. Future updates will focus on adding vector-based annotations, such as SVG, that remain editable and scalable. Additionally, plans include expanding annotation tools and exploring direct integration with data plotting libraries further to streamline the workflow from data to final figure.

## 4. Acknowledgement

We thank Nguyen Thien Luan for technical support and valuable discussion. We also thank Nguyen Thi Hai Yen, Nguyen Ky Phat, and Nguyen Tran Nam Tien for their valuable feedback.

## References

1. Clotworthy A, Davies M, Cadman TJ, Bengtsson J, Andersen TO, Kadawathagedara M, et al. Saving time and money in biomedical publishing: the case for free-format submissions with minimal requirements. BMC Medicine. 2023;21(1):172.

2. Khan A, Montenegro-Montero A, Mathelier A. Put science first and formatting later. EMBO reports. 2018;19(5):e45731.

3. Jiang Y, Lerrigo R, Ullah A, Alagappan M, Asch SM, Goodman SN, et al. The high resource impact of reformatting requirements for scientific papers. PLOS ONE. 2019;14(10):e0223976.

4. Wickham H. ggplot2: Elegant Graphics for Data Analysis. Springer-Verlag New York. ISBN 978-3-319-24277-4. 2016.

5. Kassambara A. ggpubr: ‘ggplot2’ Based Publication Ready Plots. R package version 0.6.1. 2025.

6. Pedersen TL. patchwork: The Composer of Plots. R package version 1.3.2.9000. 2025.

7. Grinberg M. Flask web development : developing web applications with Python. Sebastopol, CA: O’Reilly; 2018.

8. Feng L, Zhou N, Li Z, Fu D, Guo Y, Gao X, et al. Co-occurrence of gut microbiota dysbiosis and bile acid metabolism alteration is associated with psychological disorders in Crohn’s disease. The FASEB Journal. 2022;36(1):e22100.

9. Pang Z, Lu Y, Zhou G, Hui F, Xu L, Viau C, et al. MetaboAnalyst 6.0: towards a unified platform for metabolomics data processing, analysis and interpretation. Nucleic Acids Research. 2024;52(W1):W398–W406.

10. Lu Y, Zhou G, Ewald J, Pang Z, Shiri T, Xia J. MicrobiomeAnalyst 2.0: comprehensive statistical, functional and integrative analysis of microbiome data. Nucleic Acids Research. 2023;51(W1):W310–W8.

11. Tien NTN, Yen NTH, Phat NK, Anh NK, Thu NQ, Eunsu C, et al. Multiomics and Machine Learning Identify Immunometabolic Biomarkers for Active Tuberculosis Diagnosis Against Nontuberculous Mycobacteria and Latent Tuberculosis Infection. Journal of Proteome Research. 2025;24(8):3783–97.

12. Vosshall LB. The Glacial Pace of Scientific Publishing: Why It Hurts Everyone and What We Can Do To Fix It. The FASEB Journal. 2012;26(9):3589–93.

13. Herbert DL, Barnett AG, Clarke P, Graves N. On the time spent preparing grant proposals: an observational study of Australian researchers. BMJ Open. 2013;3(5):e002800.

14. Tien NTN, Thu NQ, Kim DH, Park S, Long NP. EasyPubPlot: A Shiny Web Application for Rapid Omics Data Exploration and Visualization. Journal of Proteome Research. 2025;24(4):2188–95.

